# Offline Replay Supports Planning: fMRI Evidence from Reward Revaluation

**DOI:** 10.1101/196758

**Authors:** Ida Momennejad, A. Ross Otto, Nathaniel D. Daw, Kenneth A. Norman

## Abstract

Making decisions in sequentially structured tasks requires integrating distally acquired information. The extensive computational cost of such integration challenges planning methods that integrate online, at decision time. Furthermore, it remains unclear whether “offline” integration during replay supports planning, and if so which memories should be replayed. Inspired by machine learning, we propose that (a) offline replay of trajectories facilitates integrating representations that guide decisions, and (b) unsigned prediction errors (uncertainty) trigger such integrative replay. We designed a 2-step revaluation task for fMRI, whereby participants needed to integrate changes in rewards with past knowledge to optimally replan decisions. As predicted, we found that (a) multi-voxel pattern evidence for off-task replay predicts subsequent replanning; (b) neural sensitivity to uncertainty predicts subsequent replay and replanning; (c) off-task hippocampus and anterior cingulate activity increase when revaluation is required. These findings elucidate how the brain leverages offline mechanisms in planning and goal-directed behavior under uncertainty.

## Introduction

To make choices in structured environments, human and non-human animals must forecast the long-term consequences of candidate actions, which often are delayed and play out serially over multiple steps. Predicting these outcomes often requires integrating information from disparate learning episodes, which occurred at different times in the past. In other words, successful planning requires both (a) accessing memories of multiple previous events and (b) integrating these memories to make useful predictions about an action’s consequences: a type of integrative memory replay (*1*). Indeed, both human and non-human animals can solve reward learning tasks designed to require this sort of integration, such as latent learning tasks (*2*) in which information about the outcome contingencies of actions (e.g., the spatial layout of a maze) is presented in a separate stage of training from the associated rewards (e.g. receipt of food).

Little is known, however, about how and when such integrative planning occurs. A longstanding assumption, dating back to Tolman (*2*) is that planning unfolds at decision time, e.g. during mental simulation of the candidate future action trajectory. More recent proposals suggest that such planning need not occur only when the animal is presented with a choice, but can also occur earlier, during *encoding* of choice-relevant information (*3*, *4*). In particular, when new information is first learned (such as food discovered in some location), its consequences for distal actions (a high value for routes leading there) can be inferred and immediately encoded. Indeed, human fMRI work provides support for both mechanisms: Studies have found that people’s ability to solve integrative evaluation problems depends on memories of distal task states accessed either at the time of choice (*5*) or encoding (*3*).

What the encoding- and choice-time mechanisms share is that both occur during active task performance, which artificial intelligence refers to as learning *online* (*6*). However, during both encoding/learning and decision-making periods, people often have limited time for exploring the set of interrelationships among decisions and outcomes, which may be prohibitive in complex environments. A third, more general possibility, which we examine here, is that integration of memories to support action evaluation can also occur *offline*, such as during off-task rest periods or even sleep. An advantage of offline planning is that it offloads computational costs to periods that are not already occupied by task behavior. This way, before the organism faces a problem that requires a decision (but after it has initially encoded the constituent events), it can update action valuations or choice policies offline, thereby accomplishing at least part of the evaluation required for later choice (*7*).

Here we investigate integrative memory processes in humans that help piece together information offline, thus supporting planning and improving decision-making behavior. In the formalism of reinforcement learning in artificial intelligence, the *Dyna* family of algorithms implements offline planning, which is accomplished by learning from *replay* of experiences that were initially encoded during online behavior (*8*) (*6*). Inspired by this work, and together with behavioral evidence of offline revaluation (*7*, *9*), computational models suggest that Dyna-like offline replay mechanisms might be implemented in the brain (*9*–*12*). Indeed, this hypothesis fits closely with a substantial rodent literature revealing that hippocampal replay of previously experienced spatial trajectories is observed during both awake behavior and sleep (*13*, *14*). It has also been shown that hippocampal replay (*15*) captures the topological structure of the task environment (*16*), and hippocampal reverse replay of trajectories occurs offline when the animal experiences changes in reward (prediction errors) at the goal state (*17*). In parallel, human fMRI research work reveals how memories are reactivated offline, such as during periods of rest that follow rewarding episodes (*18*, *19*). This offline reactivation could potentially aid in memory consolidation and integration, evidenced by enhanced subsequent memory performance (*20*–*22*). Consistent with the hypothesis that memory integration can be accomplished via offline replay, there is behavioral evidence that manipulations affecting rest periods influence humans’ ability to solve integrative decision tasks (*7*). However, so far, no animal or human study has provided neural evidence that integration during offline replay supports planning.

In this study, we use fMRI and multivariate pattern analysis (MVPA) to test the hypothesis, outlined above, that offline replay drives integrative evaluation of past policies. It has been shown that replay of a particular episode can lead to the reinstatement of neural activity in sensory cortical regions (*23*, *24*). Accordingly, we use MVPA here to track cortical reinstatement of a particular state during off-task rest periods, as a measure of offline replay of that state. We then examine the relationship between neural replay evidence (from MVPA) and subsequent choice behavior, to test whether replay facilitates the brain’s ability to integrate information and update planning policies accordingly.

The idea that replay occurs offline (rather than being triggered by particular task events) also raises the question of *prioritization*: out of all the episodes stored, which memories should the brain replay to ensure integrative and goal-directed learning? Here we draw insight again from artificial intelligence and leverage the idea of prioritized sweeping (*25*, *26*). The key proposal is that, in the face of surprising new information, memories (or states) that could lead to uncertain outcomes should be prioritized for replay. One signal of such surprise or uncertainty, which is leveraged to trigger replay in prioritized sweeping, is the experience of prediction errors—i.e. the deviation between the agent’s predictions and experiences of the world. To this end, both negative or positive prediction errors (i.e. the unsigned error) are equally meaningful as they signify the need for improved value estimates. To test this hypothesis, we use model-based fMRI analysis to evaluate whether states relating to episodes with high prediction error are preferentially replayed during subsequent rest periods.

In short, our major proposal is that that offline neural replay drives the integration of distal memories to update previously learned representations and support future planning. A further question is which memories are replayed to support planning. We suggest that episodes with large prediction errors (PEs), whether positive or negative, signal uncertainty about the optimal reward trajectory and tag related memories for future replay. This replay, in turn, enables the integration of separate memories in the service of updating optimal reward policy (action policies that maximize reward). We combine model-based fMRI, MVPA, and a sequential decision-making task to test our hypotheses.

## Results

To test the hypothesis that offline replay supports integration of memories to serve future decisions, we operationalized planning using a reward revaluation paradigm (*7*, *9*). In reward revaluation, participants first learn the multi-step sequential decisions that lead them to reward from a starting point; later, they experience a local change to later stages of the decision trajectory. Upon return to the beginning of the trajectory, we can measure the extent to which a participant’s choices adapt to the new local changes they experienced.

Here we used a 2-stage sequential decision task with a within-subject design that allowed us to manipulate whether or not revaluation was required for reward-maximizing choice behavior (*7*, *9*). In both the revaluation and control conditions, participants (*n* = 24) repeatedly visited a starting stimulus (Stage I) and chose between two actions that would lead them to one of two subsequent stimuli depending on their choice (Stage II), as displayed in Figure 1. Once a participant reached either of those Stage II stimuli (or states), they selected one of two actions to obtain reward. We will refer to the Stage I stimulus as *state 1* and the two Stage II stimuli as *state 2* and *state* 3. As a cover story, participants were told they were playing a game as a thief, exploring different buildings to steal the most money. Participants were given four separate blocks of this task, each with a different reward structure; half these blocks were randomly assigned to the *revaluation* condition, and half the blocks were assigned to the *control* condition, as described below. To ensure that participants did not transfer what they learned from one block to another, each block had a different colored background that indicated they were in a different city, and different stimuli for the Stage I and Stage II actions (Supplementary Figure 1).

**Figure 1.**
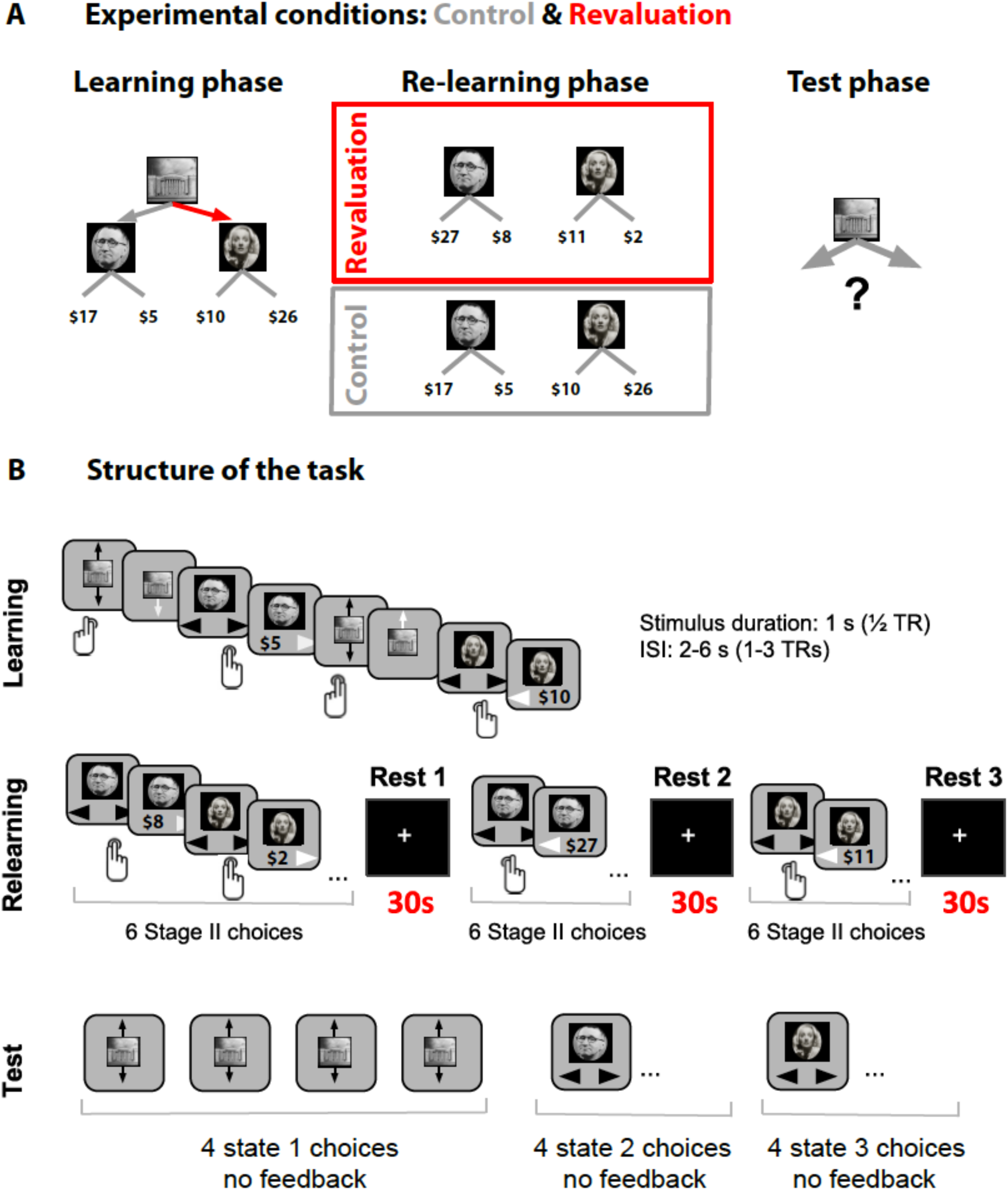
(A) Task design. Each block had 3 phases. During the Learning phase, participants explored a 2-stage environment via button presses to learn optimal reward policy. During the Relearning phase, they only explored the Stage II states: on revaluation trials the optimally rewardxing Stage II state changed during the Relearning phase; on control blocks the mean rewards did not change. The final phase was the Test phase: participants were asked to choose an action from the starting (Stage I) state that would lead to maximal reward. The red arrow denotes the action that provides access to the highest reward in the learning phase. The optimal policy during the Learning phase remains optimal in the control condition but is the suboptimal choice in the revaluation condition. **(B) The time course of an example block.**

The task in both revaluation and control blocks consisted of three phases: a Learning phase (where the participant explored the entire decision tree), a Relearning phase (in which the participant visited Stage II but never Stage I), and a Test phase (where participants revisited Stage I). During the Learning phase, participants learned the structure of states and rewards. During the Relearning phase, the rewards associated with Stage II stimuli changed for the revaluation blocks (but not control blocks). These new reward amounts in revaluation blocks were chosen so as to require a change in the decision policy leading to optimal reward in Stage I (e.g., during the Learning phase, the optimal decision policy might be to go left at Stage I, but after taking into account the Stage II reward changes experienced during Relearning, the new optimal decision policy might be to go right at Stage I). The Test phase assayed whether participants reversed their initial preference according to the Stage II changes they experienced in the Relearning phase (in control blocks we did not expect any change in preference to occur). Note that a change in Stage I policy preference would require integrating information about the action-stimulus contingencies from the Learning phase with the re-learned reward amounts associated with Stage II states from the Relearning phase. The Test phase choice was repeated four times; we computed the average proportion correct across these four choices. Importantly (to prevent any influence of new learning during the test phase), participants were not given any feedback about the consequent Stage II stimulus and reward after these choices. Note that the critical difference between the conditions was in the Relearning phase: in the control condition there was no change in the rewards associated with Stage II choices, whereas in the revaluation condition Stage II rewards changed. Hence, during the Test phase we could test whether decisions made in Stage I were affected by information learned about Stage II in the Relearning phase (see Figure 1 and Materials and Methods). As noted above, participants completed four separate blocks with this structure but different stimuli and rewards (two revaluation and two control). They did not know to which condition each block belonged (or about the logic of the manipulation), and the order of revaluation vs. control blocks was randomized.

In addition to the revaluation vs. control manipulation, we also manipulated the level of noise (variance) associated with rewards experienced following an action in a given state. This led to a 2×2 factorial design with revaluation and noise manipulations. For both revaluation and control blocks, half the blocks had variable (noisy) rewards selected from a distribution with a fixed mean (*noisy* rewards condition) leading to a persistent level of prediction errors even in the control condition (e.g. choosing left at state 2 would lead to different values that average to $5), and half of the blocks had no noise in rewards (*noiseless* rewards condition) leading to no prediction errors after learning unless there was a revaluation (e.g. choosing left at state 2 would always lead to $5). This noise manipulation—i.e. the noiseless-rewards versus noisy-rewards conditions—ensured that the difference between revaluation and control condition blocks was not limited to the absence or presence of PEs in general (i.e. a heuristic shortcut such as’any PE means revaluation’ was not possible). Put another way, the inclusion of reward noise in the noisy-rewards condition (even in control blocks) allowed us to rule out the possibility that replanning is brought about by ‘any’ experience of uncertainty, whether or not replanning was optimal. However, given that our primary interest is in replanning per se, in what follows we mainly report revaluation vs. control effects, collapsing noiseless and noisy reward conditions, while reporting statistical differences and further fMRI analyses of the noise manipulation in the Supplementary material.

Our main neural hypothesis was that offline replay would facilitate the integration of Stage I and Stage II information, which will in turn support subsequent planning behavior. Note that updating or replanning the optimal Stage I choice required integration of past information, because the transition from Stage I to Stage II, on the one hand, and the new Stage II rewards, on the other, were originally experienced at separate distal points in time during online task performance. In our study, the Relearning phase of each block was interleaved with three separate rest periods of 30s each. Thus, offline replay could be assessed using brain measurements during these off-task rest periods. To allow us to measure replay, we used image categories for Stage I and Stage II stimuli that were easy to decode with fMRI: All stimuli were either faces or scenes (*24*). In particular, the Stage I (root) stimulus belonged to a different category than the two Stage II stimuli. This manipulation allowed us to operationalize ‘offline replay’ using classifier evidence of Stage I in separate category-selective regions of interest (ROI) during rest periods, as described below.

### Replanning (revaluation) behavior

To measure revaluation behavior, we computed a *replanning magnitude* score for each subject and block (*7*, *12*, *9*), defined as the proportion of optimal Stage I choices during Test minus the proportion of choices for that same option during the Learning phase (see Equation 1 in Methods). In revaluation blocks, this replanning magnitude quantifies successful reversal of preference from Learning to Test, because the initially suboptimal option becomes optimal during the Test phase. The difference in preference captured by the replanning magnitude compares the frequency of taking the newly optimal action during the Test phase to the frequency of taking this action at the end of the Learning phase (when it was suboptimal), which adjusts for choice noisiness or failure to acquire the initial associations during the Learning phase. Thus, for revaluation blocks, a large replanning magnitude indicates consistent, successful reversal of an initially consistent, correct preference. Conversely, in control blocks (where the initially suboptimal option remains suboptimal at test) a large replanning magnitude indicates an unwarranted and suboptimal reversal of the initially trained preference, allowing us to control for preference change due to some other, nonspecific source of error such as decision noise or forgetting.

### Behavioral results

Comparison of replanning magnitudes revealed that participants showed significant revaluation behavior (i.e., reversed their Stage I choice from Learning to Test, as indicated by positive replanning magnitudes) in revaluation blocks (*t(23)=4.89, p<.0001*), but not during control blocks (*t(23)= -0.10, p=.92*), and the difference in magnitudes between the two conditions was significant (*t(23)=4.333*, *p=.0002*) (Figure 2B). The comparison with the control condition verifies that this apparent revaluation behavior is not simply attributable to general forgetting or increased memory noise from Learning to the Test phase; rather, it is due to revaluation of past policies in the face of novel information in the revaluation condition—but not the control condition. A 2-way analysis of variance confirmed the main effect of the reward vs. control factor but revealed no significant main effect of the reward stability vs. noise manipulation (see Supplementary Figure 2).

### Replay-behavior relationships

Next, we turned to the neuroimaging data to test our main hypothesis: whether neural evidence for offline replay during rest periods predicts subsequent replanning behavior. As noted above, we used MVPA to track replay of Stage I, leveraging the fact that Stage I and Stage II stimuli were always taken from different categories (faces and scenes). For example, in a trial where the Stage I stimulus was a face and Stage II stimuli were scenes, we used MVPA measures of face activity during rest periods to measure spontaneous replay of Stage I. Previous evidence suggests that multivariate pattern analysis (MVPA) can be used to measure offline replay and memory reactivation during post-task rest periods (*21*, *28*, *18*). Specifically, it has been shown that imagery, reinstatement, and perception of faces and scenes elicit patterns in the temporal cortical regions associated with visual categories, roughly corresponding to the fusiform gyrus for the face category and the parahippocampal gyrus for the scene category (*24*, *29*). Therefore, we first identified these category-selective regions using general linear models (GLMs) on a separate localizer run before the main task. The localizer run consisted of blocks of faces, scenes, objects, and rest. The faces and scenes, but not objects, were used later during the experiment. We then trained a classifier (using logistic regression) on face, scene, and object stimuli appearing during the separate localizer run. The main goal of the classifier was to classify the reinstatement of cortical patterns associated with the Stage I stimulus category during the rest periods. As a cortical measure of replay, for each participant, each block, and each of the 30s rest periods, we used this logistic regression classification method to identify TR-by-TR classifier evidence for the cortical reinstatement of faces and scenes in category ROIs (regions of interest). To ensure that we could independently assess positive evidence for both face and scene categories (rather than competitively detect one at the expense of the other), we assessed classifier evidence for each category from an ROI that was selective for that category (see Methods for further discussion of this point).

To test whether offline replay during intermittent rest periods in the Relearning phase predicted successful revaluation in the later Test phase, we averaged MVPA evidence for reinstatement of the Stage I category during all rest periods (Figure 2, *x*-axis), and computed the correlation with mean revaluation behavior in each condition (Figure 2, *y*-axis), across subjects. Even though the Stage I stimulus category never appeared during the Relearning phase, it was important to ensure that we were not simply observing residual signals evoked by the Stage II stimuli observed prior to each rest phase. Therefore, we excluded the first 5 TRs (corresponding to 10 seconds) of the beginning of each rest period from analysis.

In revaluation trials, there was a significant correlation between MVPA evidence for Stage I replay during the three off-task rest periods and revaluation behavior during the subsequent Test phase (*rho=0.54, p=0.0068*) but not for control trials (*rho=-0.13, p=0.55*, Figure 2). The correlation coefficients for the revaluation vs. the control trials were significantly different from one another (*p=.0230*; bootstrap, see Methods). We also conducted the same analyses, separately, for the noisy rewards versus the noiseless rewards conditions. The revaluation vs. control difference was trending in the noisy condition (*p =.066*) but not the noiseless condition (*p = .16*), and there was no significant interaction between revaluation/control and noisy/noiseless (*p = .34*) (Supplementary Figure 2; see Discussion section for further treatment of these results).

**Figure 2.**
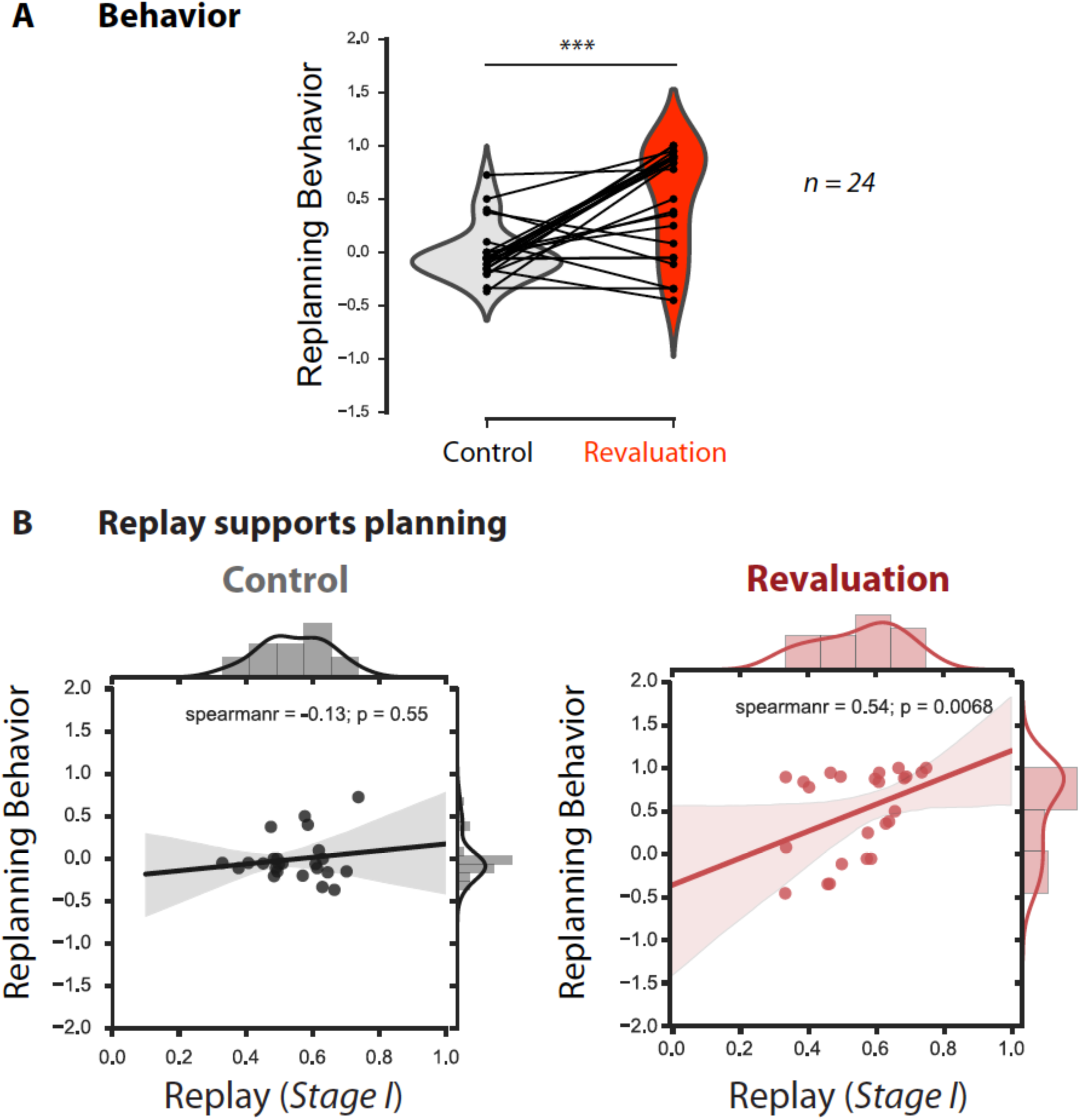
Offline replay of distal past states supports planning. (A) Behavioral results (n=24). Participants significantly changed their choice from the Learning phase to the Test phase in the revaluation condition, but not the control condition. **(B) Replay supports planning.** Correlation between Stage I replay during off-task rest periods and replanning magnitude during the subsequent Test phase for Control (left) and Revaluation (right) blocks. Stage I replay was operationalized as MVPA evidence for the Stage I stimulus category, in category-selective regions of interest, during all rest periods of control and revaluation conditions (n=24). The correlation was significant in the Revaluation blocks (Spearman rho=.54, p=.0068) but not Control blocks (Spearman rho=-0.13, p=0.55), and it was significantly larger in Revaluation than Control blocks (p=.0230, computed using a bootstrap, computing Spearman rho 1000 times with replacement). Regression lines are provided for visualization purposes, but statistics were done on Spearman rho values.

In short, we found that MVPA evidence of Stage I replay during rest correlates with subsequent replanning behavior. This fits with our hypothesis that offline memory processes can support planning. Next, we test the hypothesis that this replay is driven by the experience of prediction error.

### The brain’s response to unsigned prediction error tags memories for replay

A key question for any account of learning from replay is how the brain selects which memories to replay. We hypothesized that the brain’s sensitivity to episodes with surprising information or increased uncertainty, here operationalized as neural sensitivity to unsigned PEs (i.e., the absolute value of PEs), would ‘tag’ the memory of states and trajectories related to that episode with priority (*25*) for later offline replay. This subsequent offline replay in turn would support updating past policies and hence predict future revaluation behavior (Figure 3, note the relationships indicated in the diagram, top left). We have shown in previous sections that offline replay during rest is correlated with revaluation during a later Test phase. To examine evidence for the hypothesized relationship between the brain’s response to uncertainty and offline replay, as well as subsequent replanning behavior (Figure 3, top left), we ran parametric modulation analyses as follows.

For each participant, we conducted a parametric regression analysis within the Revaluation condition to identify regions with sensitivity to stimulus-by-stimulus unsigned PEs. Prediction errors were estimated from a Temporal Difference (TD) learning model (see Methods). We then tested, for each subject and each Revaluation block, the relationship between the magnitude of the brain’s PE response (in that block) and two different covariates: subsequent replay and subsequent replanning score (Figure 3, top right, purple and blue maps correspondingly). This allowed us to determine whether (and in which brain areas) the magnitude of activity in response to unsigned PE was related to off-line replay (Figure 3, regions indicated in purple, see Supplementary Table 1 for coordinates) and subsequent replanning (Figure 3, regions indicated in Blue, see Supplementary Table 2 for coordinates), as hypothesized (Figure 3, top left). In order to identify the combined effect and given the correlation between the two covariates, we used a conjunction analysis over the two resulting correlation brain maps (Figure 3, regions indicated in green). This conjunction analysis yielded regions where the brain’s response to unsigned PE on each block correlated with both (a) subsequent mean replay evidence (for Stage I stimuli) during rest, and (b) subsequent replanning magnitude of the choice behavior during that block’s Test phase (Figure 3).

**Figure 3.**
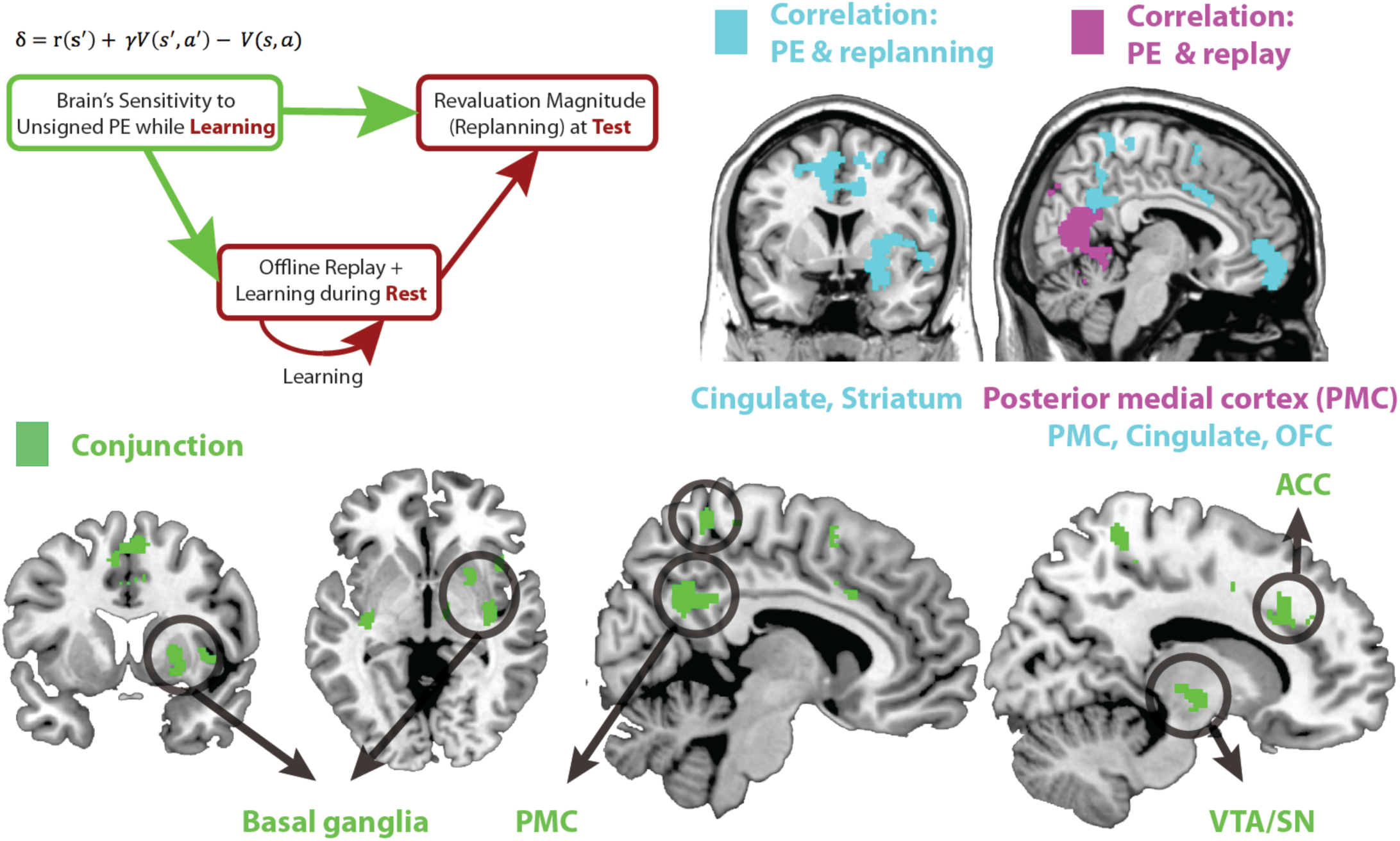
(Top left) Schematic of a theoretical account of revaluation. We propose that neural sensitivity to reward prediction errors during learning ‘tags’ or ‘prioritizes’ memories for replay during later rest periods. Replay during rest, in turn, allows the comparison of past policy with new simulated policy and updating of the past policy when needed. **(Top right)** Regions where sensitivity to unsigned PE in Revaluation blocks correlates with subsequent replay during rest (extent threshold p<.005, cluster family-wise error [FWE] corrected, p<.05, shown in purple) and revaluation behavior (extent threshold p<.005, cluster FWE corrected, p<.05, shown in blue). **(Bottom)** Green reveals the conjunction of regions where sensitivity to unsigned PE in Revaluation blocks correlates with subsequent replay during rest (extent threshold p<.005, cluster FWE corrected, p<.05) and revaluation behavior (threshold p<.005, cluster FWE corrected, p<.05) in those blocks; the conjunction is shown at a p<.05 threshold. We found that the sensitivity of broad regions in the basal ganglia, the cingulate cortex (including the ACC), and the posterior medial cortex (precuneus) to unsigned prediction errors (signaling increase in uncertainty) correlated with both future replay during rest as well as subsequent revaluation behavior. See Supplementary Tables 1 and 2 for coordinates.

This conjunction analysis yielded multiple regions where neural sensitivity to unsigned PEs significantly predicted both future offline replay and revaluation magnitudes: the anterior cingulate cortex (ACC), the mid cingulate, posterior medial cortex (PMC) including dorsal and ventral precuneus, and the basal ganglia, especially the putamen (Figure 3, bottom). These results are consistent with the hypothesized relationship between sensitivity to prediction errors during learning, offline replay during rest, and revaluation behavior at test, and suggest an involvement of ACC, PMC, and basal ganglia in signaling the relevant PEs (see Supplementary Figure 3 for breakdown of this analysis for the noisy-rewards and noiseless-rewards blocks).

### Brain activity during revaluation vs. control rest

Our hypothesis is that learning and updating take place during offline replay periods—indeed, we revealed above that replay in category-sensitive brain regions during rest periods of revaluation blocks (but not control blocks) predicted subsequent planning behavior. This hypothesis also implies that brain regions related to learning and memory should show more activity during the rest periods of revaluation trials, where engaging in offline replay could result in better choices, compared to control trials in which the most learning has already occurred during the Learning phase. To test this, we used a GLM to compare overall differences in univariate activity during the rest periods of revaluation vs. control trials. Contrasting rest periods from revaluation trials versus control trials (Rest _revaluation>_ Rest _control_) revealed higher activity in the hippocampus/medial temporal lobe and the anterior cingulate cortex (p<.005, cluster corrected at p<.05, Figure 4, see Supplementary Table 3 for coordinates, and Supplementary Figure 4 for breakdown of this analysis for the noisy-rewards and noiseless-rewards blocks).

### Learning during rest

So far, we have shown that evidence for offline replay, across subjects and blocks, correlated with subsequent replanning behavior, and that regions broadly involved in memory and evaluation were more active during the rest period of revaluation trials. Finally, we broke down these effects over the course of the three rest periods in the Relearning phase to examine how they evolved over the progression of relearning and seek evidence of learning dynamics.

A property of the learning model is that PEs during the Relearning period should decrease from the period preceding Rest 1 (when participants are first encountering the new rewards in the revaluation condition) to the period preceding Rest 3 (after participants have had more time to learn these new rewards); this was the case for the PE values estimated by the learning model (Figure 5, top left). Given the decreasing pattern in unsigned PE during re-learning, we predicted that this within-block variation might be echoed in patterns of replay across rest periods – reduced PE should lead to less replay. To focus the analysis on differences among rest periods within each block, we mean-centered MVPA evidence for the Stage I stimulus during the 3 rest periods for each block of each subject. This allowed us to compare patterns of within-block differences between replay in the three rest periods of revaluation and control trials. In control blocks, we found no differences in MVPA evidence for S1 replay during the three rest periods (Figure 5). During revaluation blocks, however, evidence for offline replay significantly decreased from Rest 1 to Rest 3 (*t(23)=2.9, p=.007*), similar to the decreasing pattern of unsigned prediction errors (Figure 5). Furthermore, offline replay was higher during Rest 1 of revaluation blocks compared to the Rest 1 of control blocks (*t(23)=2.17, p=.039*). Thus, consistent with our hypothesis, prediction errors and replay track each other over the course of the block, as well as across blocks (as previously shown in Figure 3).

**Figure 4.**
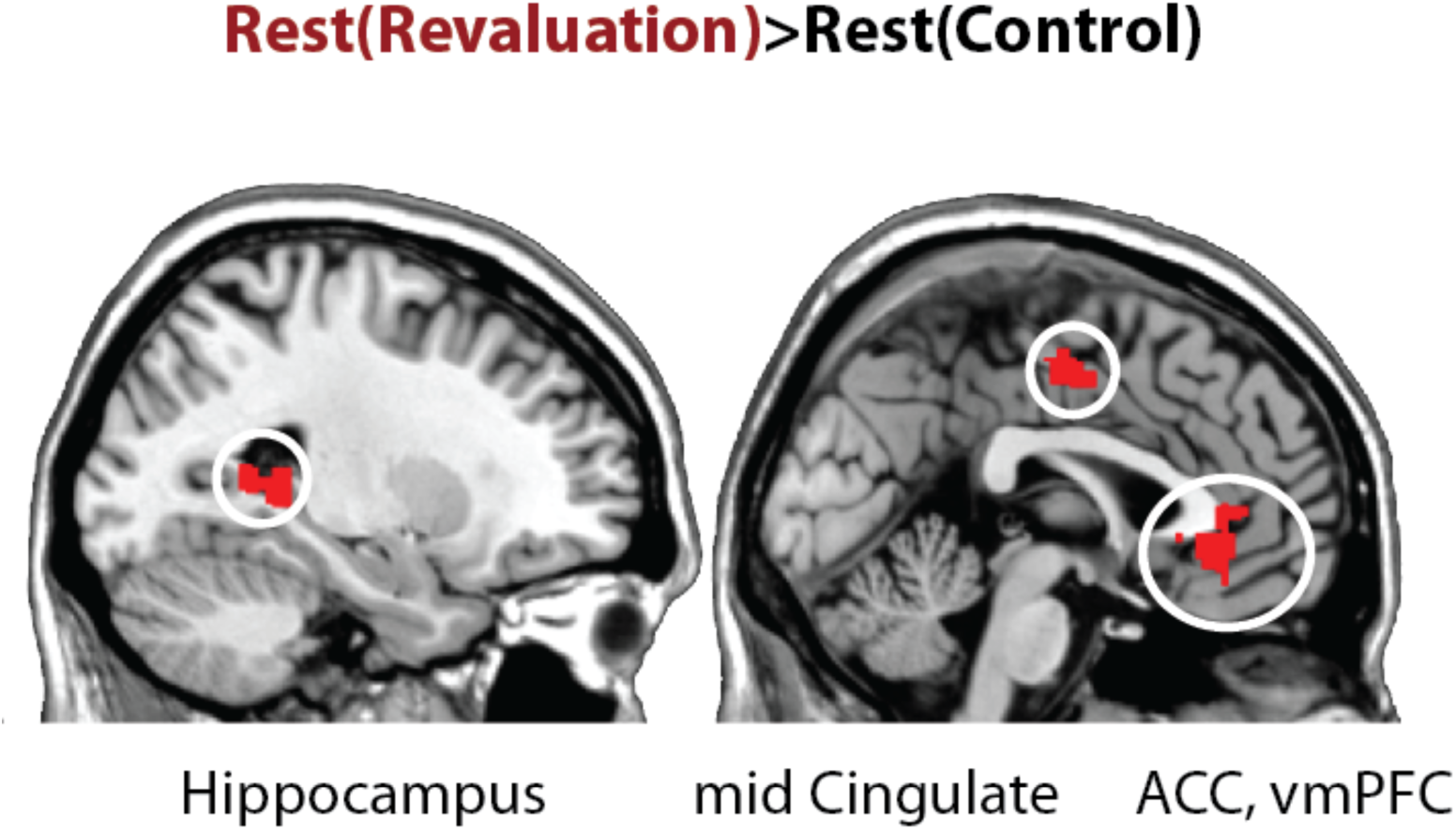
Univariate general linear model contrasts. Comparing activity rest periods of revaluation vs. control blocks, Rest_revaluation>_ Rest_control_. The contrast reveals higher activity in the hippocampus, the anterior cingulate cortex, mid cingulate (shown above), as well as bilateral insula and superior temporal cortices (extent threshold p<.005, cluster level family-wise error corrected at p<.05). See Supplementary Table 3 for coordinates, cluster size, and p^values^.

**Figure 5.**
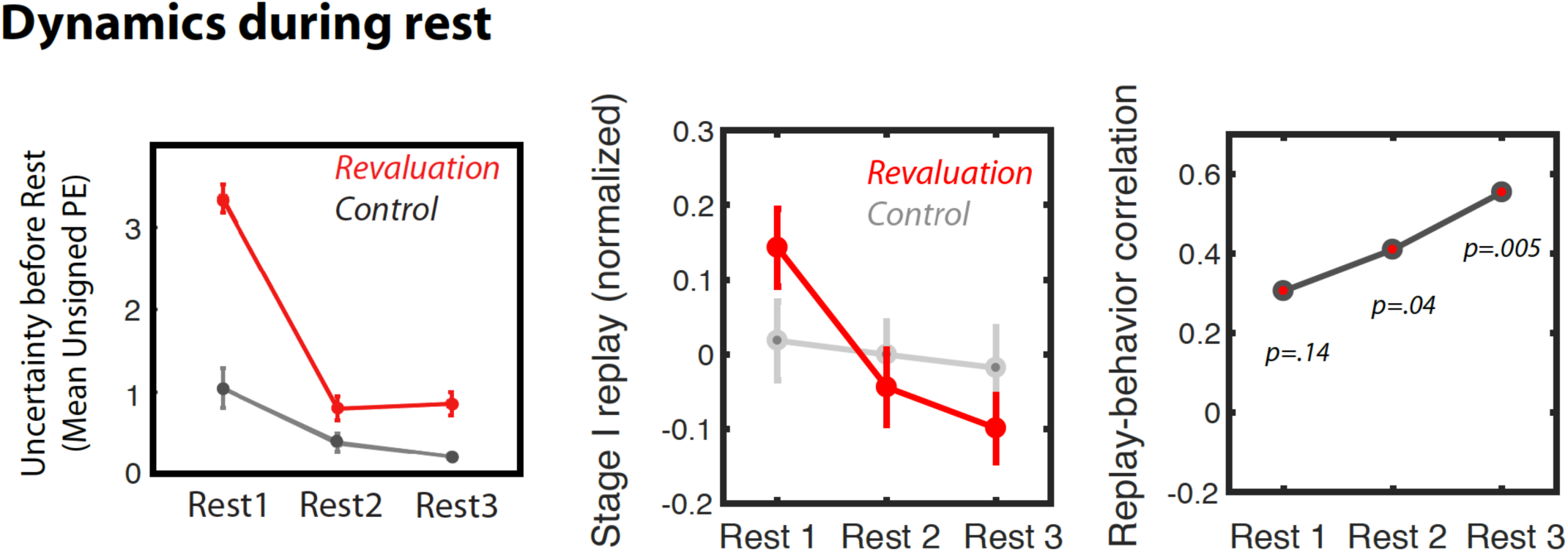
Dynamics of replay and activity during rest periods. Dynamics of prediction errors prior to each rest period (left), MVPA evidence for Stage I replay during each rest period (middle), and replay-behavior correlation across the 3 rest periods of revaluation runs (right).

Analyzing the rest periods of revaluation trials in the noisy-rewards and noiseless-rewards conditions separately (Supplementary Figure 5), we found a similar trend: In the noisy-rewards condition, replay was marginally higher during Rest 1 than both Rest 2 (*t(23)=2.07, p=.049*) and Rest 3 (*t(23)=2.02, p=.05*) – in the noiseless-rewards condition, replay levels were low overall and did not vary across rest periods.

Finally, we considered how this replay related to subsequent replanning behavior in the Test phase. We compared the replay-replanning correlation across the different rest periods (Figure 5; unpacking over time the correlation from Figure 2B). We found that, for the average of all revaluation blocks, this correlation was numerically stronger (and only statistically significant) during later rest periods (however, there was no significant difference between the correlation coefficient for Rest 1 versus the correlation coefficient for Rest 3, Fisher’s *z=-1.01, p=.31*). We then analyzed this effect separately for revaluation blocks in the noisy rewards and revaluation blocks in the noiseless-rewards condition (Supplementary Figure 5), and found again that the replay-behavior correlation was mainly driven by the noise condition, where the most significant replay-behavior correlation is observed during Rest 2.

## Discussion

We present fMRI evidence for the hypothesis that uncertainty-driven replay can help integrate past episodes offline, and in turn, update representations and policies that support planning (*8*). First, we found that MVPA evidence for offline replay correlates with subsequent planning behavior. We suggest that during offline replay the brain integrates and pieces together simulated trajectories of information acquired during multiple learning episodes and updates planning policies accordingly. Second, we hypothesized that prediction errors drive offline replay; in support of this, we found that the brain’s sensitivity to unsigned prediction errors in the ACC, basal ganglia, and posterior medial cortex (the precuneus) predicted the extent of subsequent offline replay as well as ensuing replanning behavior. We suggest that the brain’s sensitivity to surprising information and increased uncertainty, here signaled by unsigned prediction errors, may tag certain episodes and trajectories of experience for selective offline replay later on (*25*, *26*). Finally, we observed a general increase in hippocampal, posterior medial, anterior cingulate (extending to ventromedial prefrontal cortex upon relaxing the threshold), and superior temporal cortex activity during the rest periods of revaluation trials compared to control. The hippocampus has long been identified as the locus of episodic memory (*30*), posterior medial cortex is shown to be involved in recall and integration of memories over long time-scales (*31*, *32*), the anterior cingulate cortex is known to enable comparisons and signal conflicts the resolution of which may be valuable enough to recruit controlled processes (*33*), and the ventromedial prefrontal cortex has been established as a region involved in memory retrieval of relevant information (*34*, *35*), value representation (*36*), and decision making (*37*) respectively. Hippocampal-prefrontal interactions have been shown to mediate control of memory in memory inhibition (*38*) and updating memory in extinction (*39*). Thus, this finding supports an increased role for memory and evaluation mechanisms during the rest periods of revaluation blocks.

In short, inspired by work on the Dyna architecture in reinforcement learning (*6*) and prioritized sweeping (*25*, *26*) in machine learning, as well as behavioral paradigms of reward revaluation (*7*, *9*), these results are the first to show a functional role for offline replay in planning under uncertainty. The key novel aspect of our results is our finding that offline replay can lead to replanning by *integrating multiple, temporally distal learning episodes* (here, computing the implication of Stage II experiences on choice policy at Stage I). Previous human studies have shown benefits of *online* reinstatement of past episodes on decision-making in a sensory preconditioning paradigm (*3*), and that goal-directed behavior in sequential decision-making is accompanied by the online reinstatement of prospective states (*5*). Other studies have explored properties of offline reinstatement, showing that it favors past episodes associated with higher reward (*18*) and supports upcoming learning of related material (*28*). However, to our knowledge, no human or rodent study has tested the role of offline replay in piecing together distal episodes to support planning. Tolman famously designed revaluation tasks that require the animal to integrate information experienced at various points in time (*2*). Thus, we used a simplified Tolman-style revaluation task to test the role of offline replay in integration of distant memories and address the crucial question of which memories the brain would preferentially replay to improve planning. While the rodent literature has shown in various paradigms that offline replay is associated with better task performance (*10*, *14*, *17*), none verifies that these improvements are specifically due to the offline integration of memory. Our present findings further these previous efforts and elucidate the role of offline replay in goal-directed memory integration that supports planning.

Given that our interest is primarily in the relationships between prediction errors, replay, and revaluation behavior, we focused our analysis on the difference between revaluation and control conditions, collapsing over blocks with noisy vs. noiseless rewards. We did, however, also consider all results separately across those conditions. Perhaps unsurprisingly (given lower power), we did not find significant differences between the conditions in any of our measures, but we note a few findings. First, we only observed a significant correlation between replay and replanning behavior in revaluation blocks in the noisy reward condition (Supplementary Figure 2). These results may be due to a ceiling effect in the noiseless reward conditions, where updating past policies was sufficiently easy for online processes to adequately do the job. Second, we only found significant conjunction results for the PE analyses in the combined condition, but neither in the noisy-rewards condition nor the noiseless-rewards conditions alone. This is most likely due to the fact that the data in the noisy and noiseless conditions were half the number of runs included in the combined analysis reported in the main text (Figure 3). Future experiments should focus on the differential effect of the noisiness vs. stability of reward structures, and the volatility of event and reward structures more broadly, on brain regions recruited in offline processes that support planning. One possibility is that the brain is more likely to benefit from offline replay for planning when the reward structure in the world is noisier, in which case changes in mean reward may be neglected as variance, making online integration more difficult. While the present study has focused on reward prediction errors, future studies are required to study the role of other forms of prediction errors, e.g. state prediction errors (*40*).

We also examined the dynamics of our results over the progression of the relearning phase. The key difference between the three rest periods was in the amount of unsigned prediction error (PE, or uncertainty) experienced prior to each (Figure 5). Here, unsigned prediction errors are how the brain notices something is not right, i.e. some prediction does not match the observation, and needs to determine whether the surprising experience may have further consequences for the brain’s stored representations and policies (its model of the world). Before Rest 1 during revaluation trials, participants had just noticed that rewards have changed from what they expected, leading to the experience of high unsigned PEs while making Stage II decisions. Importantly, they did not visit the Stage I state during the Relearning phase, but only Stage II stimuli, therefore they did not experience any PEs after choosing a policy from Stage I. Before Rest 3 of revaluation trials, participants experienced smaller prediction errors because they had already had the chance to learn the new reward structures for Stage II stimuli; correspondingly, overall levels of replay were lower after Rest 3 than after Rest 1. However, crucially, while levels of replay were lower after Rest 3, there was still a robust correlation between Rest 3 replay and behavior (the size of the replay-behavior correlation did not significantly differ across rest periods).

Taken together, these findings suggest a potential two-stage functional role for replay in the present study. First, replay of trajectories marked with unsigned PE could allow identifying distal parts of the state space that may need to be updated. When the brain encounters prediction errors, initial replay can enable search through a graph of memory states to identify other states affected by the experienced PE: this is the “sample to tag with priority’ function of replay. The present experiment consists of one such past state, but it is possible that in larger decision trees this search for relevant past states leads to ‘reverse replay’ (*16*, *17*). As such key past states that lead to the present state in which high PE was experienced may be tagged with priority over other memories to be replayed later on. Second, once the new structure of the environment is adequately learned, and high PEs are no longer experienced; the tagged relevant past states and their corresponding trajectories can be replayed, simulated if never directly experienced, and updated. A similar tagging idea had been proposed in the domain of fear learning, where memory traces could be retroactively altered in accordance with new behavioral relevance of fear associations (*41*), whereas here we propose a more general case where unsigned PEs tag memories with priority for subsequent integrative replay. As such, the second function of replay may be ‘learning from replay’: to actually enable the brain to piece together and simulate trajectories in order to update past representations and planning policies offline (*15*).

It is worth commenting on how the notion of replay here relates to replay in the rodent literature – in this literature, the term ‘replay’ often refers to the replay of sequences at sub-second speed. The temporal resolution of fMRI poses a challenge to providing direct neural evidence of sequential replay. That said, we designed the study such that replanning the optimal Stage I choice required integration of past information by piecing together different memories sequentially: The transition from Stage I to Stage II, on the one hand, and the new Stage II rewards, on the other, were originally experienced at separate distal points in time during online task performance. Unlike animal studies, but similar to previous human fMRI studies of offline replay (*21*, *28*, *18*), here we leverage a controlled design together with evidence of Stage I reinstatement to investigate the role of offline replay in planning. Future work using methods with higher temporal resolution, such as MEG and direct recordings from the hippocampus in patients, is required to shed light on the details of forward and reverse sequential replay in updating representations and memories that serve planning.

In future work, we plan to conduct more detailed explorations of the functional role of replay in planning, using paradigms with larger decision trees and higher temporal resolutions that make it possible to track replay trajectories in more detail; e.g., a recent human study used MEG to identify a role for reverse replay of trajectories (*42*). We also plan to explore how offline replay might contribute to generalization and temporal abstraction, e.g. via multi-step predictive representations of upcoming states (or the successor representation, (*12*, *9*). Caching of multi-step predictive representations during replay can in turn enable generalization (*43*), faster planning especially under time pressure, and sub-goal discovery (*44*) across longer scales. After consolidation, the generalized multi-step representations could lead to abstract representations of states and policies such as schemas. Using generalized representations, both discovery of relevant past states (as in the first function of replay above) and piecing together episodes to simulate trajectories to goal (as the second function of replay above) could also take place in a multi-step fashion consistent with the successor representation (*12*), enabling more efficiency simulation of past and future trajectories.

To summarize, we have provided neural and behavioral evidence for the hypothesis that offline replay supports planning. We further suggest that the brain’s sensitivity to uncertainty, operationalized as unsigned prediction errors, mediates the amount of offline replay and, through this, revaluation behavior. Our findings further our understanding of how the brain leverages offline memory mechanisms in planning and goal-directed behavior.

## Material and Methods

### Subjects

We recruited 26 volunteers using the Princeton University recruitment system (SONA) to participate in the fMRI study. Two participants were excluded due to running out of time and not having sufficient number of runs. The remaining 24 participants” data were further analyzed. The Princeton University Institutional Review Board approved the study. All participants gave informed consent to participate in the fMRI study and signed a screening form that ensured they had normal or corrected to normal vision, had no metal in their body, and had no history of psychiatric or neurological disorders.

### Stimuli and task

Eight face, eight scene, and eight object stimuli were used in the face/scene/object functional localizers. The faces and scenes were subsequently used in the main experimental task. Before the beginning of the main experiment, participants were subjected to blocked functional localizer runs. In the localizer participants were exposed to separate blocks of scenes, faces, and objects. Each category was presented for 4 blocks that each lasted for 12 s, separated by rest periods of 12 s. In each block, every stimulus was presented for 3.5 s with an ISI of .5 s. To ensure that participants were paying attention to the stimuli during the localizer, they were instructed to give responses to a cover task. In half of the localizer blocks participants were instructed to give responses with the left hand and in the other half of the blocks with the right hand.

The main experimental paradigm consisted of 4 runs, each of which corresponded to one Markov Decision Process (MDP) with three phases. Two of 4 runs were assigned to the revaluation condition and the rest were control runs (in which no revaluation took place). The order of conditions was randomized for each participant. The task was explained to the participants in terms of a cover task with a robbery scenario: They were told the experiment was a game where they were to explore different locations in different cities, each corresponding to each run’s simple 2-step maze to find out which state they could steal more money from. Participants were instructed that they would receive a bonus compensation for their performance on the final Test phase, which they received for all runs at the end of the fMRI session. Each MDP was presented as a new ‘city’ in which the participants would explore a building in search of money. Different cities were randomly assigned to different conditions across participants. Each MDP consisted of 3 main states: Stage I and two Stage II states. The same stimulus signaled Stage I in each city, and the same Stage II stimuli signaled states 2 and 3. Importantly, Stage I and Stage II stimuli belonged to different categories. In 2 out of 4 cities state 1 was a face and states 2 and 3 were scenes, and in the remaining two cities the opposite was the case.

Additionally, we manipulated the variance in the rewards observed at each state. We wanted to ensure that, after learning, participants did not only experience prediction errors in the revaluation condition, but that the control condition also contained some degree of uncertainty with respect to rewards. To this end, the rewards in half of the runs, i.e. one revaluation run and one control run, were sampled from a normal distribution with a fixed mean and variance, and in the other half of runs the rewards were completely deterministic.

In each city, during the Learning phase participants were exposed to the stimuli associated with the three main states of the MDP about 70 times. The last 10 choices they made from the Stage I state were used to calculate their probability of choosing the optimal response (e.g. if the optimal policy was left: p(left) = #left choices/10). Following the Learning phase, participants entered the Relearning phase where they only visited Stage II stimuli and never the Stage I stimulus. During this phase in revaluation trials, the rewards associated with Stage II (state 2 and 3) choices changed – but not during control trials. After every six Stage II episodes each requiring a choice, a rest period of 30 s length followed for which no instructions were given. Each participant experienced three rest periods of 30 s each during the Relearning phase. Finally, during the Test phase, participants were once again exposed to state 1, four times. Replanning magnitudes were computed as the probability of choosing the other valid action in those four trials, minus the probability of choosing the same response during the Learning phase – thus detecting the actual change in performance by subtracting noisiness in decision at the end of the Learning phase. For instance, if the optimal policy at test was right: revaluation magnitude = (#right choices at test/4) - (#right during last 10 choices of learning/10).

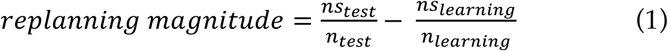

Here *n*_*test*_ denotes the total number of valid responses (i.e., excluding missed responses or invalid button presses) during the Test phase, and *ns*_*test*_ is the number of times that the optimal choice was made during the test phase; *n*_*learning*_ denotes the total number of valid responses during the last 10 trials of the Learning phase,and *ns*_*learning*_ is the number of times that the choice that was optimal at Test was selected during the last 10 trials of the learning phase (note that, in revaluation blocks, the response that is optimal during the Test phase was suboptimal during the Learning phase).

### fMRI imaging protocol and preprocessing

Imaging data were acquired on a 3 Tesla Skyra scanner (Siemens, Erlangen, Germany) using a 20-channel head coil. Functional runs were acquired using T2^*^-weighted echo-planar sequence with 37 slices (interleaved order, ascending) with 3-mm thickness, an in-plane resolution of 3×3 m, TR=2.08 s, echo time of 70 ms, anterior-to-posterior phase encoding direction, and no gaps. Slices were tilted for each participant by roughly 30 degrees to avoid the sinuses and optimize signal in the orbitofrontal cortex but avoid losing temporal cortex voxels. A T1-weighted structural volume was acquired with 1×1×1 mm resolution. Functional volumes were slice-time corrected, realigned to the mean of each run, and motion corrected. FMRI data processing was carried out using FEAT (FMRI Expert Analysis Tool) Version 6.00, part of FSL (FMRIB’s Software Library, http://www.fmrib.ox.ac.uk/fsl). Higher-level analysis was carried out using FLAME (FMRIB’s Local Analysis of Mixed Effects) stage 1 (*45*–*47*).

### Behavioral and neuroimaging data analysis

#### Functional localization

We designed a general linear model (GLM) to identify functional regions for each participant. We used regressors corresponding to face, scene, objects, and rest conditions, and applied Scene>Face and Face>Scene contrasts to identify face-selective and scene-selective regions for each participant (*29*); for these contrasts, we used a liberal voxel-wise threshold of p < .005, uncorrected, to ensure that we did not miss informative voxels. These regions of interest were used as a means of dimensionality reduction, allowing us to confine the multivariate analysis of experimental runs to these regions.

##### Multivariate Pattern Analysis

As noted, Stage I stimuli were of a different category than Stage II stimuli. For each participant, Stage I stimuli were faces in half of the blocks and scenes in the other half. This ensured that identifying Stage I evidence was not confounded with simply a replay bias for faces and scenes, which might have been the case if the Stage I stimuli were from the same category across all trials. We used L2-regularized logistic regression (penalty=1) to train a classifier on TRs when participants observed images of faces, objects, and scenes during the independent functional localizer run acquired prior to experiment (*24*). Importantly, to detect face activation, we trained a face vs. other (scene, object) classifier only on voxels from the face-selective ROI identified in the localizer analysis above; conversely, to detect scene activation, we trained a scene vs. other (face, object) classifier only on voxels from the scene-selective ROI identified in the localizer analysis above. Our use of this approach (where we used separate sets of voxels for face detection and scene detection) was driven by our desire to obtain, to the greatest extent possible, *independent* readouts of face and scene activity. It is a well known issue with MVPA that, when face and scene classifiers are applied to the same voxels, face and scene classifier readouts tend to be artifactually anti-correlated because the face and scene labels are anti-correlated in the training data (i.e., the classifier learns that, in the training data, the presence of scene predicts the absence of face, and vice-versa) (*48*). This problem can be mitigated to some degree by adding additional categories at training (this is why we added the object category), but we found that it can be mitigated even further by applying the classifiers to distinct sets of voxels – this carries a potential cost of losing sensitivity (by restricting the voxels going into a particular classifier) but we were willing to pay this price to address the anti-correlation problem described above: In our study, the correlation between the time course of face evidence (from face ROIs) and scene evidence (from scene ROIs) during the rest periods was *positive* (on average, .36), not negative.

To measure replay of Stage I category information, we took the trained face and scene classifiers and applied them to the TR-by-TR volumes from each run’s rest periods, focusing specifically on the last 10 TRs of each rest period (we omitted the first 5 TRs from each rest period to minimize “spillover” from participants perceiving Stage II stimuli right before the rest period started). As noted above, if the Stage I stimulus was a face, we used the face classifier to detect replay; if the Stage I stimulus was a scene, we used the scene classifier. For each TR, the classifier outputs an evidence value for the selected category between zero and one; we averaged these evidence values across TRs to get a neural replay score for each block (since there were 3 rest periods per block and we used 10 TRs per rest period, the replay score for each block was based on 30 TRs).

The main goal of this analysis was to test whether evidence for offline replay of the root state (State 1) during the rest periods correlated with subsequent revaluation behavior during the Test phase. Therefore, we computed correlations between mean classification accuracy (evidence for offline replay of Stage I) and subsequent revaluation behavior separately for revaluation condition trials and control trials (Figure 2B).

##### Bootstrap comparison of replay-replanning correlation across conditions

To compare the size of the correlation between replay and behavior across conditions (revaluation vs. control), we used a bootstrap approach in which we resampled participants with replacement 1000 times; for each bootstrap sample, we computed the difference in the size of the replay-behavior correlation across conditions, thereby giving us a bootstrap distribution of correlation differences.

### Univariate analysis of rest periods

#### Model-based analysis

In order to identify regions that were sensitive to reward prediction errors (RPE), we ran a parametric modulation analysis in which we regressed out mean signal change when participants received reward, and identified regions where variance around the mean was modulated by the magnitude of absolute RPEs experienced while receiving the reward. Unsigned prediction errors were calculated for each subject and each stimulus using equation (2).

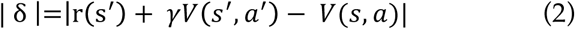

Here *s* refers to the current state and *s*’to the next state, *a* is the current action and *a*’ the next action, and γ is the temporal discounting parameter (here γ= 1, since the temporal horizon in a 2-step Markov decision process (MDP) is limited to one step (*40*)). The reward prediction error is in turn used to update the state-action value as equation (3) below, where α denotes the learning rate. We estimated a learning rate of .7 based on analysis of previous pilot behavioral data (n=24) and used this learning rate for all participants.

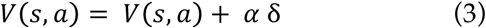

We were interested in identifying regions where sensitivity to unsigned reward PEs, a measure of uncertainty used in the reinforcement learning literature, would correlate with future replay during rest periods and revaluation magnitude during the final Test phase.

## Acknowledgments

We thank Matthew Botvinick and Sam Gershman for helpful conversations, and Sarah Dubrow, Dylan Rich, and James Antony for helpful comments on manuscript drafts. We acknowledge and thank Eeh Pyoung Rhee, who ran control analyses to compare classifier performance for his senior thesis in computer science at Princeton University. This project and publication was made possible through the support of a grant from the John Templeton Foundation, grant 57876, and NIMH grant R01MH109177, part of the CRCNS program. The opinions expressed in this publication are those of the authors and do not necessarily reflect the views of the John Templeton Foundation or other funding bodies.

---

**Supplementary Figure 1.**
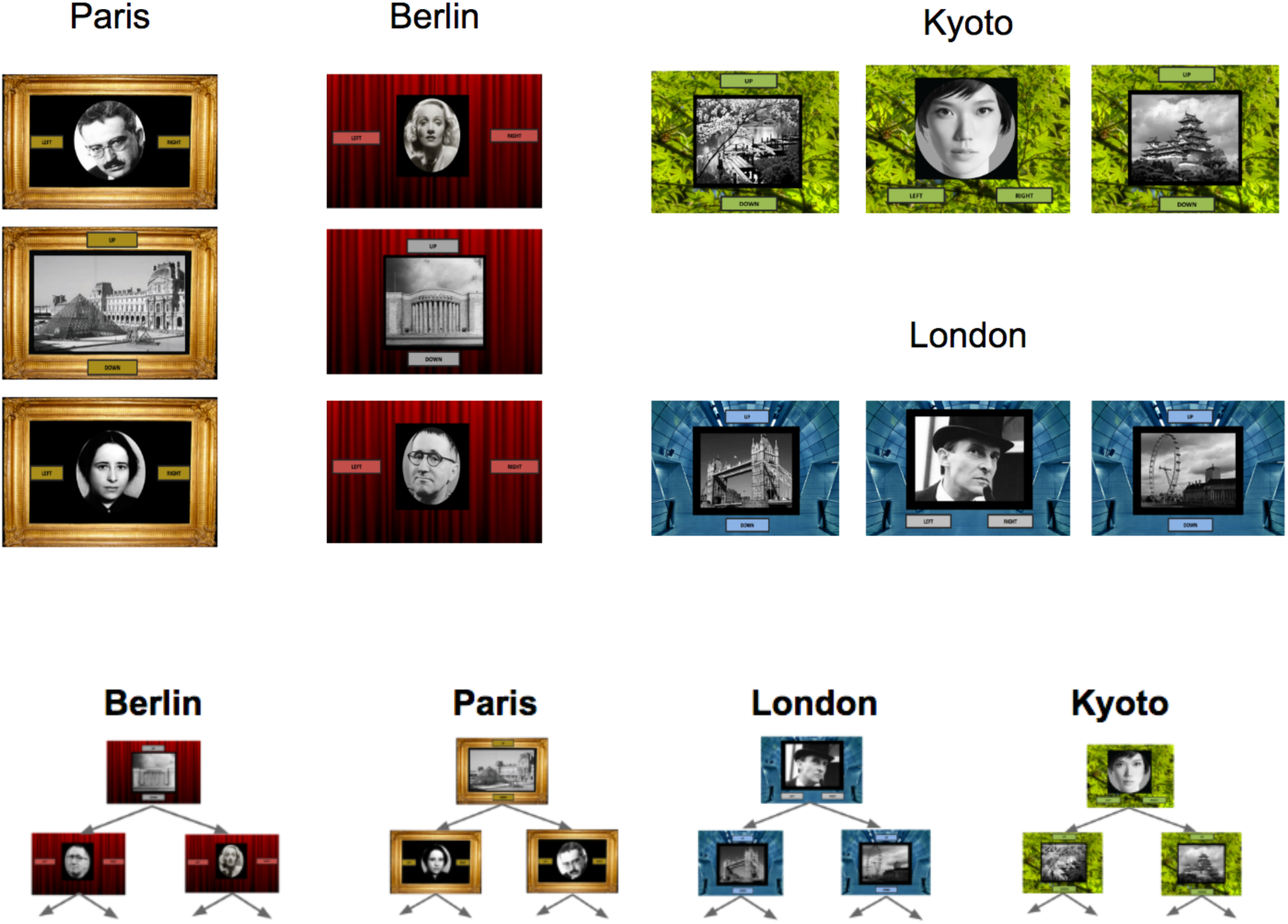
The four different blocks. The cover story described each block, with its own Markov decision process (MDP), in terms of stealing money in a new building in a new city. For each participant, half of the cities were used in the revaluation condition and half in the control condition. In each of these conditions, one block had noiseless rewards (participants experienced the same reward each time they visited a state) and one block had noisy rewards (participants experienced the reward of a given state with variance around the fixed mean reward).

**Supplementary Figure 2.**
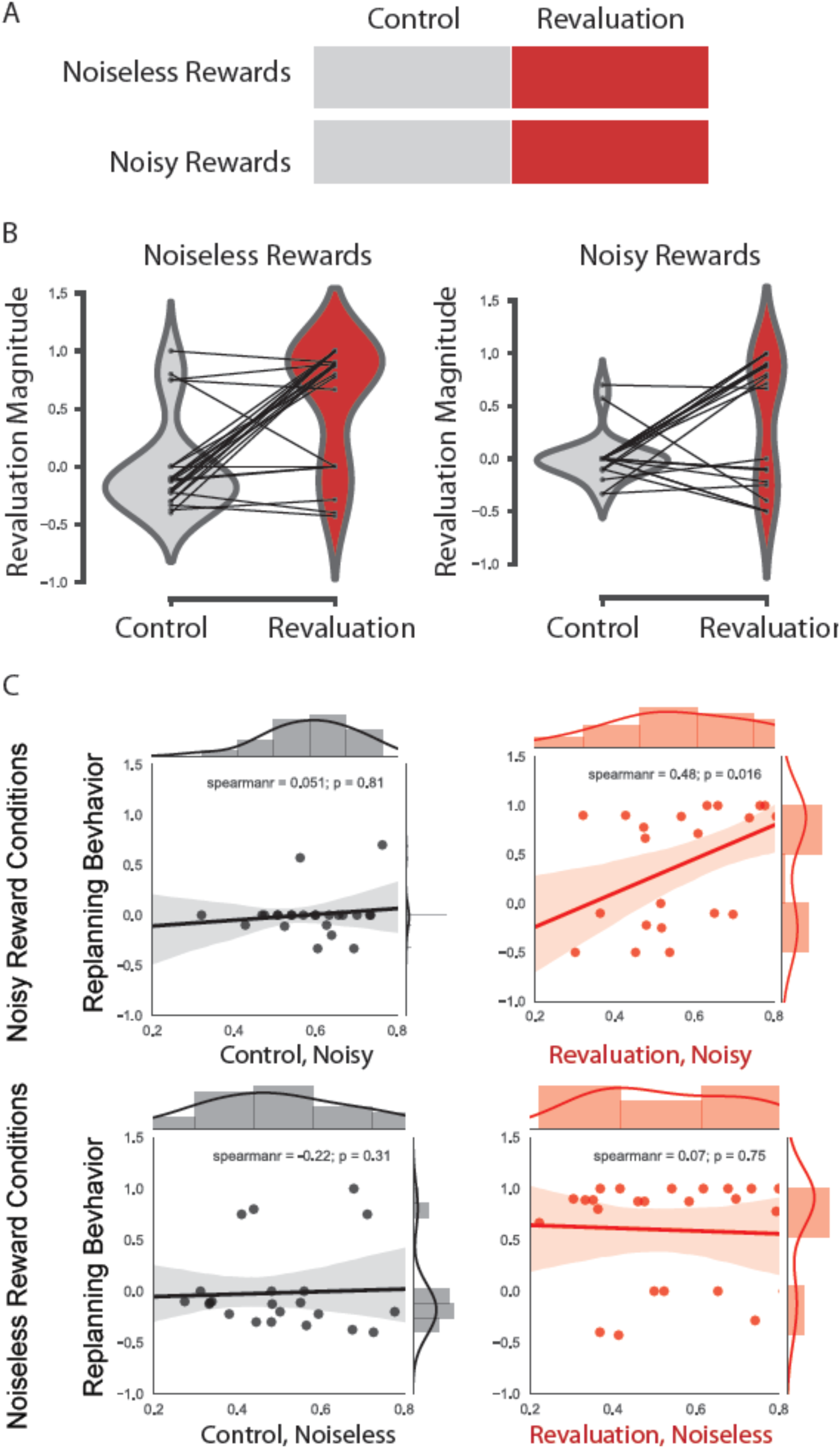
(A) Illustration of experimental conditions obtained by crossing revaluation and noise. (B) Behavioral revaluation scores in the control and revaluation conditions, assessed separately in the noisy rewards vs. noiseless rewards conditions. Half of the trials in the control and revaluation conditions had fixed rewards (noiseless rewards condition) and half had noisy rewards (noisy rewards condition). We used an ANOVA to compare replanning behavior, evidenced by the revaluation magnitude, in the conditions with no variance in the rewards (noiseless condition) vs. conditions with noisy reward (noisy condition). Analysis of variance revealed a significant effect of the revaluation condition on behavior (F(1, 23)=29.57, p<.0001) but no significant effect of the noise condition (F(1, 23)=.91, p=.34), and no significant interactions (F(1, 23)=1.35, p=.25. Within each of the noise conditions (noiseless, noisy), participants significantly changed their choice from the Learning phase to the Test phase in the revaluation condition, but not the control condition: noiseless rewards condition: t(23)=4.6, p=.00003; noisy rewards condition: t(23)=3.06, p=.003. **(C) Offline replay of distant past states predicts replanning**. Breakdown of correlation between MVPA evidence for replay and subsequent replanning behavior, computed separately for the noisy and noiseless rewards conditions. All correlations were conducted using the last 10 TRs (out of 15 TRs) of each rest period, excluding the first 5 TRs to reduce residual effects of Stage II stimulus-presentation prior to rest. This breakdown revealed that MVPA evidence for Stage I replay was significantly correlated with subsequent replanning behavior in revaluation blocks in the noisy rewards condition (Spearman’s rho = .48, p = .016), but not the noiseless rewards condition (Spearman’s rho = .07, p = .75). We then ran bootstrap analyses to assess the differences in correlations between conditions. The difference in correlations between revaluation and control was marginally significant in the noisy rewards condition (p = .066) but not in the noiseless condition (p = .16). There was no overall interaction between revaluation/control and noisy/noiseless; i.e., the revaluation vs. control difference was not significantly larger in the noisy condition than the noiseless condition (p = .34).

**Supplementary Figure 3.**
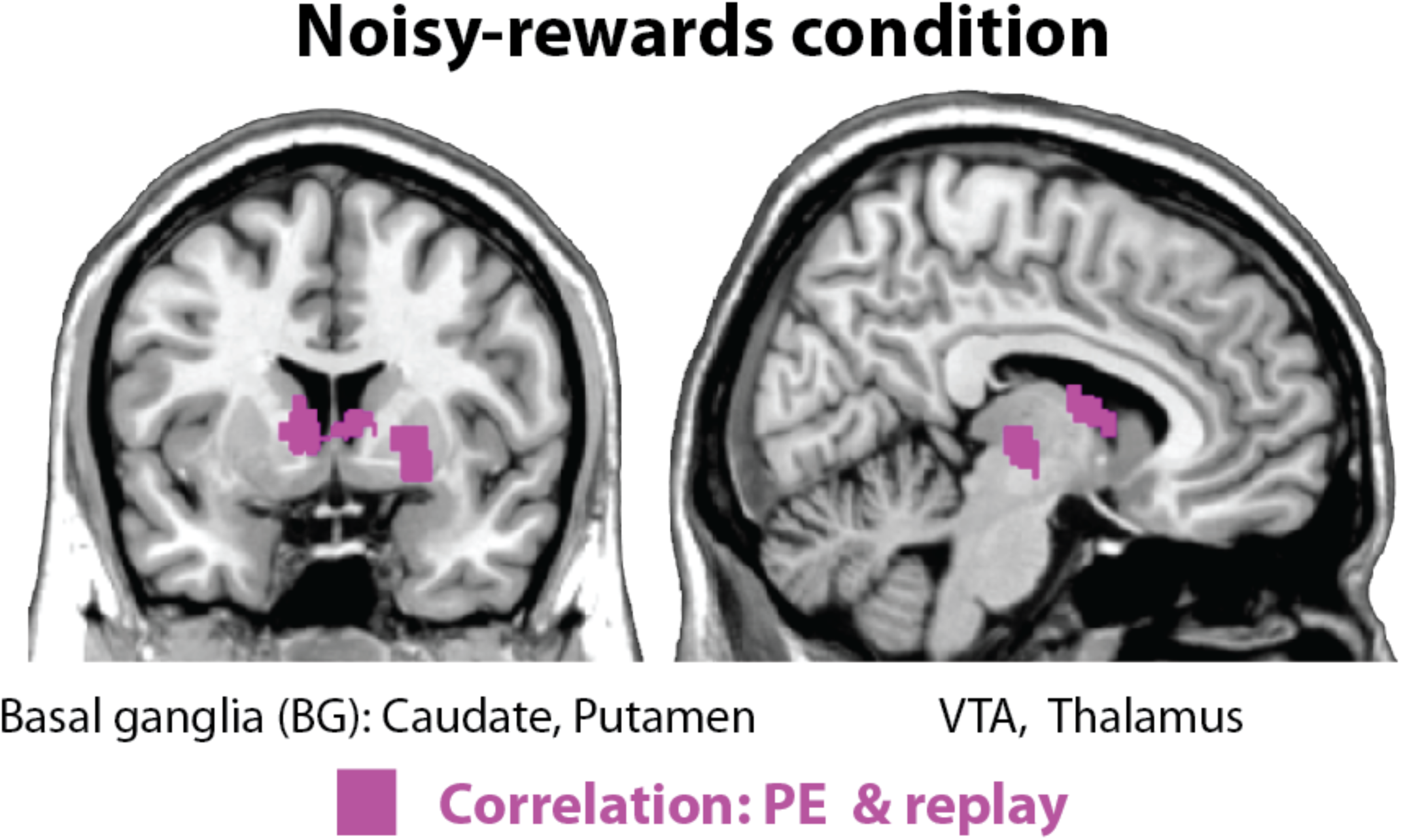
Correlation between the brain’s response to unsigned prediction errors and subsequent replay (purple), replanning behavior (blue), and their conjunction (green) in revaluation blocks, run separately for the noisy and noiseless conditions. Results are shown based on extent threshold p<.005 and cluster (family wise error) correction at p<.05. The only one of these analyses to yield any significant clusters was the correlation between PE and replay in the noisy condition (shown below), in which the magnitude of sensitivity to unsigned PE in the ventral tegmental area (VTA), the thalamus, and the basal ganglia (including the caudate and putamen) predicted subsequent replay during rest periods.

**Supplementary Figure 4.**
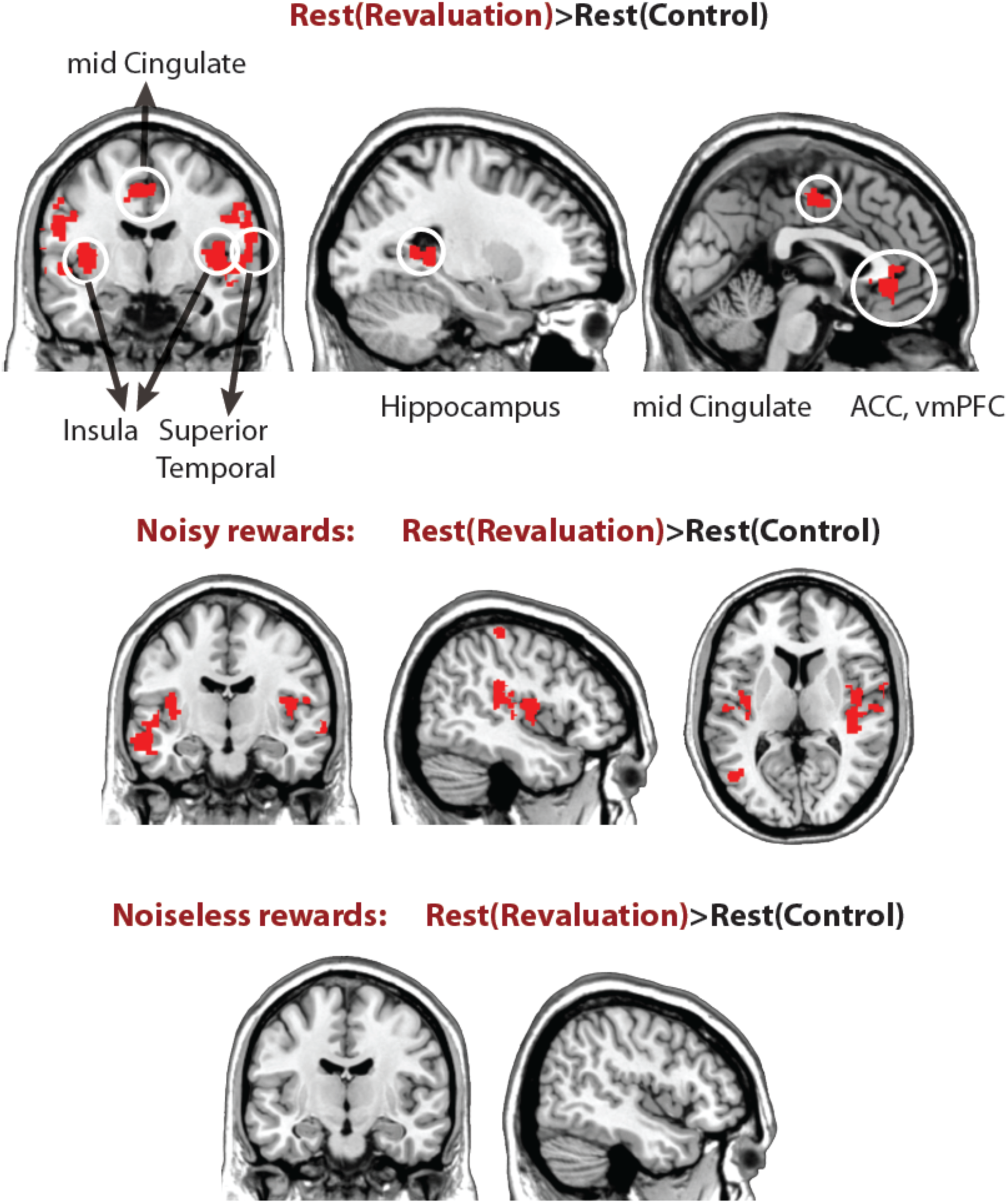
Differences in off-task univariate activation in revaluation vs. control. We compared the difference between univariate activation during the rest periods of control vs. revaluation blocks in all runs (top), in the noisy-rewards condition (middle), and the noiseless-rewards condition (bottom). The Rest(revaluation)>Rest(control) contrast in the noisy rewards conditions reveals lateral temporal cortices and the insula, whereas the noiseless condition did not reveal any significant regions (extent threshold p<.005, cluster FWE corrected, p<.05).

**Supplementary Figure 5.**
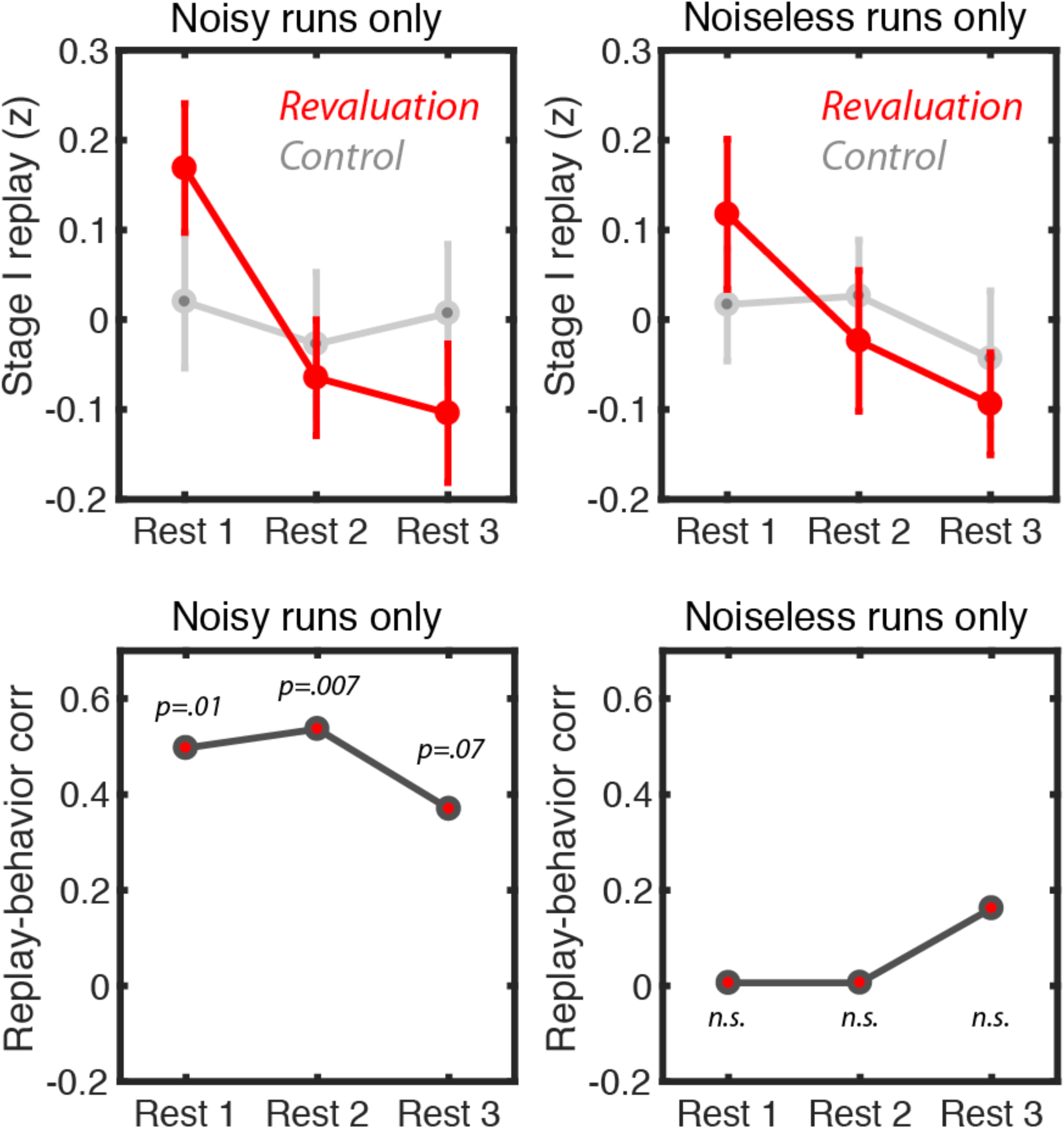
Breakdown of replay during the three rest periods in the revaluation vs. control conditions during noiseless rewards and noisy rewards conditions (top) and the correlation of these replay magnitudes with revaluation magnitude (bottom) in the noiseless vs. noisy rewards conditions.

**Supplementary Table 1.** Coordinates of voxels where parametric modulation with unsigned PEs during learning predicted future replay of Stage I during rest periods, extent threshold p<.005, corrected at cluster level family-wise error p<.05 (these correspond to purple regions in Figure 3).

| **Region** | **Z score** | **x** | **y** | **z** | **k voxels** | **p** |
| --- | --- | --- | --- | --- | --- | --- |
| *Right Cuneus* | 3.31 | -2 | -68 | 4 | 1898 | .000488 |
| *Left Cuneus* | 3.29 | 4 | -66 | 22 |  |  |
| *Right Lingual* | 3.21 | -14 | -68 | 22 |  |  |
| *Left Lingual* | 3.15 | 16 | -46 | -8 |  |  |
| *Calcarine* | 3.14 | 6 | -70 | 4 |  |  |

**Supplementary Table 2.** Coordinates of voxels where parametric modulation with unsigned PEs during learning predicted future replanning behavior (revaluation magnitude), extent threshold p<.005, corrected at cluster level family-wise error p<.05 (these correspond to blue regions in Figure 3).

| **Region** | **Z score** | **x** | **y** | **z** | **k voxels** | **p** |
| --- | --- | --- | --- | --- | --- | --- |
| *Right precentral* | 3.67 | 26 | -20 | 68 | 5240 | 1.36e-08 |
| *Orbitofrontal cortex (OFC)* | 3.62 | 0 | 58 | -12 |  |  |
| *Supplementary motor area* | 3.59 | -8 | 4 | 48 |  |  |
| *Right superior frontal cortex* | 3.49 | 18 | 32 | 38 |  |  |
| *ACC* | 3.49 | -14 | 50 | 0 |  |  |
| *Right OFC* | 3.42 | 6 | 58 | -18 |  |  |
| *Right superior temporal* | 3.67 | 56 | -30 | 14 | 5165 | 1.67e-08 |
| *Right supra-marginal* | 3.6 | 34 | -36 | 44 |  |  |
| *Inferior parietal* | 3.58 | 36 | -42 | 44 |  |  |
| *Right putamen* | 3.39 | 26 | 8 | 2 |  |  |
| *Right superior temporal pole* | 3.35 | 46 | 18 | -16 |  |  |
| *Left superior temporal pole* | 3.4 | -46 | -10 | 0 | 2161 | .00021 |
| *Left putamen* | 3.35 | -20 | 18 | 8 |  |  |

**Supplementary Table 3.**
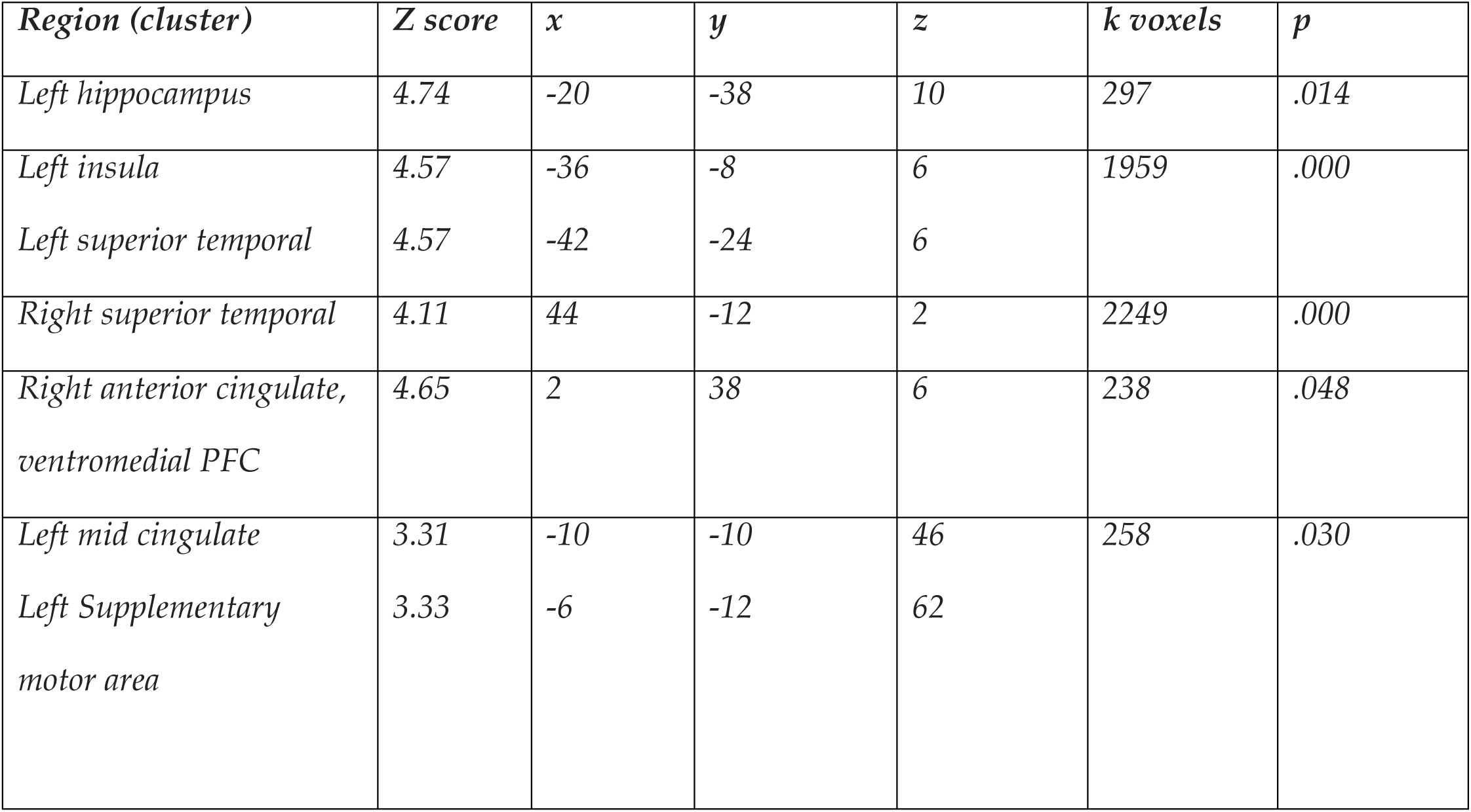
Coordinates of peak voxels of regions with higher off-task activity during rest periods of revaluation>control condition. The clusters were selected with threshold p<.005, corrected at cluster level family-wise error p<.05 (these correspond to red regions in Figure 4).

